# Prior selection prevents the loss of an ecosystem cycle during acidification

**DOI:** 10.1101/2020.01.27.921437

**Authors:** Sofia J. van Moorsel, Justin N. Marleau, Jorge O. Negrín Dastis, Charles Bazerghi, Vincent Fugère, Owen L. Petchey, Andrew Gonzalez

**Author notes:** E-mail addresses for correspondence.

## Abstract

Ecosystem processes vary temporally due to variation in environmental variables, such as when diurnal variation in sunlight causes diurnal cycles in net primary production. This variability can be characterized by its frequency and amplitude, used to define “normal” functioning of an ecosystem. Relatively little research has addressed how normal modes of variability, such as diurnal cycles, are lost or recovered, following anthropogenic stress. We conducted an aquatic mesocosm experiment to test whether prior application of environmental stress, in the form of moderate acidification, affected the diurnal cycle of dissolved oxygen when exposed to severe acidification. High-frequency data from sensor loggers deployed in 12 mesocosms showed that severe acidification caused a temporary loss of diurnal variation in dissolved oxygen concentration. However, pre-exposure to an acidic environment resulted in the persistence of the diurnal cycle. We hypothesize that pre-exposure shifted the community to acid tolerant genotypes and/or species of algae and other photosynthetic organisms. Our findings suggest that the stability of ecosystem cycles is intrinsically liked to the stress tolerance of the species assemblage.

## INTRODUCTION

Anthropogenic sources of environmental stress such as acidification affect the structure and function of ecosystems by impacting their geochemistry and species composition. Stressors can drive the loss of intolerant species and depending on the degree of compositional turnover, incur a suite of different destabilizing effects such as a loss of diversity (Geelen and Leuven 1986, Niyogi et al. 2003) and/or keystone species (Cuenca Cambronero et al. 2018) and a synchronization of its constituent populations (Thompson et al. 2015). A community that has experienced a stressor may differ in structure, function, and stability (e.g. variability stability, or resistance) to those that have not experienced the stress (Ives and Cardinale 2004, Keitt 2008).

Ecosystem processes are known to vary over a range of time scales. Some scales of variability arise from periodic responses to natural cycles in the environment, while others are scales reflect the aggregate response of the community to stochastic variation in the environment (Blasius et al. 1999, Gilg 2003, Keitt 2008). Because scale-specific responses to the environment occur at different periods and amplitudes, it is necessary to assess ecosystem stability at different time scales (Downing et al. 2008). Cyclical responses to natural environmental changes depend on the time scale in focus, from daily (night/day), seasonal (length of day) and annual (length of growing season) fluctuations. For example, freshwater ecosystems are characterized by strong daily fluctuations in oxygen production (Schindler et al. 2017). During the day, when light is present, algal production of oxygen via photosynthesis can be greater than all organismal consumption of oxygen via respiration, leading to increase in dissolved oxygen concentration. In contrast, at night, in the absence of photosynthesis, organismal respiration decreases dissolved oxygen. The higher the primary productivity, the stronger these diurnal cycles (Schindler et al. 2017).

The growth of stress tolerant species can recover ecosystem processes from a certain degree of stress (Rapport et al. 1985). However, at doses initially lethal for all species only rapid evolution can allow the recovery of an exposed community (unless the system is open to immigration of resistant genotypes). Previous work has shown that exposure to nonlethal stress can pre-adapt populations to a subsequent exposure to otherwise lethal press perturbation allowing resident species and genotypes to adjust their physiology and demographic rates (Bell and Gonzalez 2009, 2011, Low-Décarie et al. 2015, Fugère et al. 2020). Exposure to a stressor may thus reduce the effects of subsequent exposure to the same stressor (Bell and Gonzalez 2009, 2011, Low-Décarie et al. 2015, Fugère et al. 2020) due to selection for tolerant/resistant genotypes within the populations comprising the community.

The diurnal cycle of DO is a metric for ecosystem functioning (Venkiteswaran et al. 2008, Demars et al. 2015, Schindler et al. 2017) capturing the activity of aquatic species across trophic levels from decomposers to zooplankton (Cowan et al. 1996). We therefore tested the hypothesis that a pre-selective environment could enable communities to maintain their diurnal DO cycle when subsequently exposed to a lethal level of stress, i.e. maintain a ‘normal’ (as measured in controls) frequency and magnitude of diurnal variability of the DO cycle during the press acidification treatment. We take a “black-box” approach with this whole-ecosystem approach, which comes at the expense of specific mechanisms. We used a high-resolution DO time series from a 5-month long experiment in mesocosms containing 1000 liters of lake water and natural plankton communities. We carried out a multi-phase experiment using acidification as the stressor. In phase 1, we imposed selective environments by manipulating pre-exposure to sublethal stress (with weekly addition of sulfuric acid to the ponds). Then, in phase 2, all ponds were exposed to a one-time lethal dose of sulfuric acid resulting in pH 3 continuously for several weeks.

## METHODS

### Field site and design

This mesocosm study was conducted within the Large Experimental Array of Ponds (LEAP) platform at the Gault Nature Reserve in Mont-St-Hilaire, QC, Canada (45°32’ N, 73°08’ W, 122 m a.s.l.). The experiment was run between May and October 2018 for a total of 147 days. On 24 May 2018, 105 mesocosms (1100L stock tanks, Rubbermaid, Huntersville, NC, USA), henceforth referred to as ponds, were filled with approximately 1000 liters of unfiltered lake water from nearby oligotrophic Lac Hertel, located 1 km upstream of the experimental facility. Lac Hertel is situated within a UNESCO biosphere reserve and has a fully forested and protected watershed, free of agricultural run-off and other pollution. The lake has no recorded history of acidification. One day after the ponds were filled, all ponds received 50 ml of a nutrient solution containing nitrogen and phosphorus (86.128 g/l KNO_3_, 1.78 g/l KH_2_PO_4_, 2.24 g/l K_2_HPO_4_, which resulted in the total addition of 596.45 g of N and 39.8 g of P) to increase nutrient levels and stimulate primary production.

As a measure of ecosystem functioning, we used dissolved oxygen (DO). DO is both determined by biological factors (e.g. community biomass and composition) and by environmental factors such as water temperature (Belley et al. 2016), as the solubility of oxygen decreases as temperature increases (Wetzel 2001). We tracked DO dynamics in twelve ponds using sensor loggers (MiniDOTs, PME, Vista, California, USA) attached to the side of the ponds with rope and positioned into the water column at a depth of approximately 20 cm. We focus our analysis on this subset of 12 mesocosms that generated high frequency time series of DO. Water temperature (°C) and dissolved oxygen (in mg/l) was measured every 20 minutes. Ponds were covered with 1 mm netting (vegetable garden netting) to prevent insects, foliage, and debris from entry. Periphyton growth on the side of the ponds or on the loggers was minimal. Four loggers were in mesocosms at pH 5.5, four at pH 6.5 and four at pH 8.5 (see Fig. 1). We present dissolved oxygen corrected for temperature as % saturation in Fig 1 and all subsequent analyses (described below) focused on temperature-corrected values (% saturation). Outlier values > 4 standard deviations away from the mean DO across all data points (98 %) were attributed to temporary probe disturbance or malfunction and were thus excluded from analyses of DO across time and mean DO per phase. These values (DO saturation < 40 % or > 150%) represented 0.4 % of all data points. Due to the robustness of the wavelet analysis, no data points were excluded from those analyses. We assumed that all ponds had a similar wind exposure and gas exchange coefficient, and thus did not correct oxygen data for gas exchange with the atmosphere.

**Figure 1.**
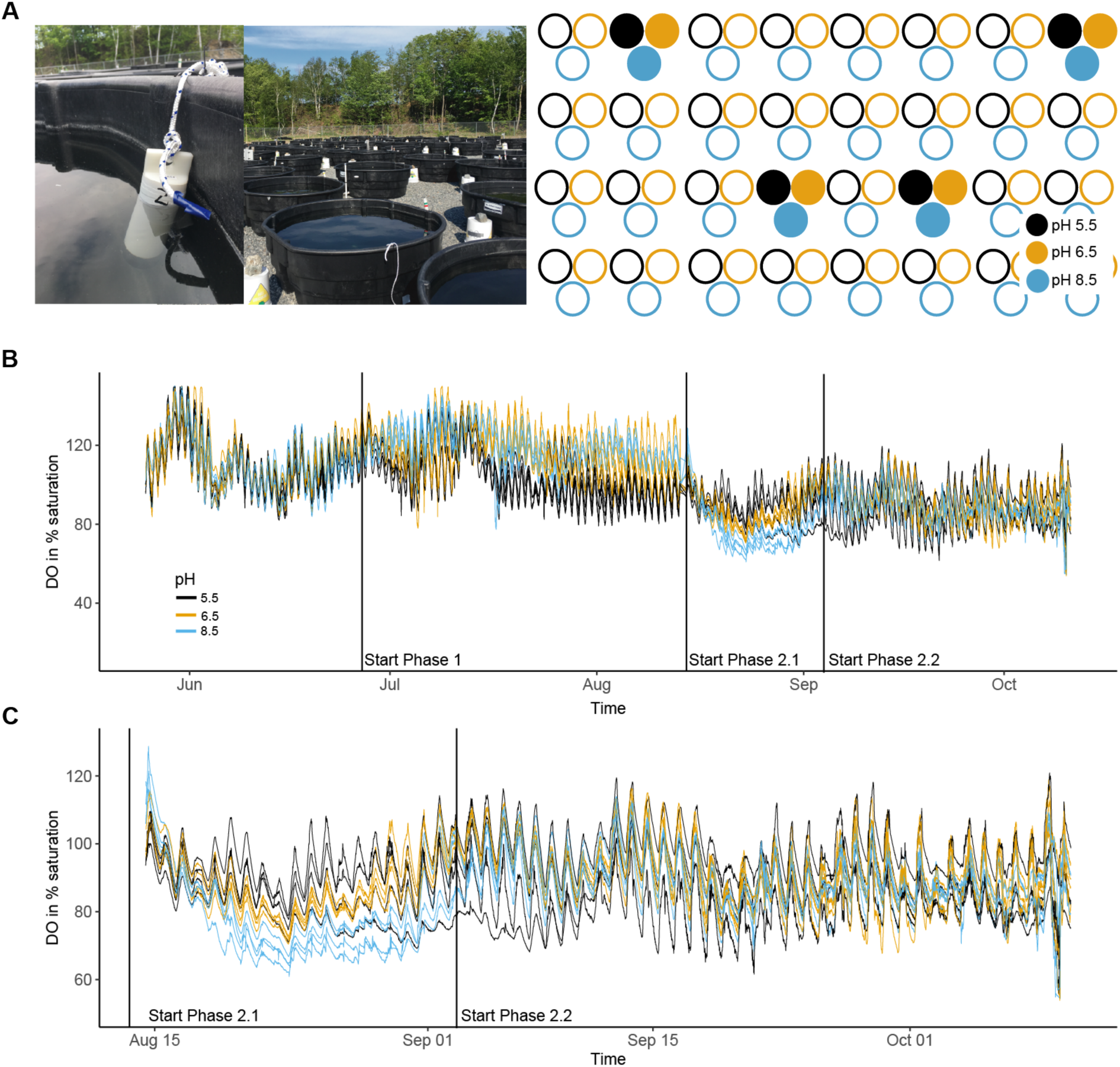
**(A)** Photographs of the field site and spatial arrangement of the experimental ponds. Filled circles indicate the ponds in which the miniDOT sensor loggers were deployed. **(B)** Dissolved oxygen in % saturation over time according to pH treatment during phase 1. **(C)** Close up of phase 2 for DO in % saturation. The start of phase 1 and 2 are indicated with a dashed vertical line. Black lines, pH 5.5, orange lines, pH 6.5 and blue lines, pH 8.5.

### Acidification treatment

During the four weeks of phase 0 from 24 May to 26 June ecosystems were at their natural pH (phase 0, mean pH = 9.03± 0.17 on June 7, 8.27± 0.15 on June 12 and 8.81± 0.26 on June 26, Fig. S2). We acidified the pond to induce selection on community composition. Acidification is harmful for freshwater systems causing population declines for higher trophic levels (e.g. fish and zooplankton) and shifts towards a simplification of community composition for lower trophic levels, such as phytoplankton (Locke and Sprules 1994, Schindler et al. 1996, Weiss et al. 2018). On day 34 of the experiment (26 June 2018), we acidified the ponds to start our selection treatment (phase 1). Using sulfuric acid, four ponds were acidified to pH 5.5 and four ponds were acidified to pH 6.5. The remaining four ponds were left as they were at pH 8.5. Sulfuric acid was gradually added to the ponds using a pipette and stirred. pH was measured using a multiparameter sonde (YSI, Yellow Springs, OH, USA). Weekly acidification maintained the pH around the target pH-value. Initially strong buffering capacity resulted in weekly increases of the pH, but we succeeded in establishing three distinct pH levels during phase 1 (Figure S2).

On day 83 of the experiment (14 August 2018), we started phase 2 of the experiment. All ponds were acidified using sulfuric acid to pH 3, except pond A4 which was accidentally acidified to pH 2.5 because the pH sensor malfunctioned (see Figure S2). On August 16, we acidified again to ensure that all ponds were +/- 0.1 from pH 3; pH remained stable thereafter until the end of the experiment. The pH in Pond A4 never recovered and remained at pH 2.5 so we conducted our analyses with and without this pond for comparison (Figure S2).

### Wavelet analysis

Under normal conditions the ponds exhibit a daily cycle in DO due to daily cycles in the amount of photosynthesis occurring, hence we were interested in identifying if this daily cycle persisted throughout the acidification treatments across the phases of the experiment. We were also interested in whether the degree of persistence and cycle amplitude depended on the acid pre-exposure treatment. To detect change in the daily period and its amplitude over time we used two continuous wavelet transform algorithms (Grinsted et al. 2004) in Matlab (v2018a) that allowed us to both detect oscillations at multiple frequencies as well as locate their occurrence in time. For our main results, we used the Morlet wavelet (Fig. 2; Grinsted et al. 2004) and the Morse wavelet (Fig. 3; Lilly and Olhede 2012), which lead to similar scalograms, but different normalization schemes, indicating robust results (see Supplementary Information for more details). To identify whether peaks in variability at particular frequencies were statistically significant we used autocorrelated noise as the null variation (Grinsted et al. 2004). With the Morse wavelet transform algorithm, we could extract the amplitude of the daily oscillations for each pond over each phase of the experiment by taking the modulus (or absolute value) of the complex coefficients of the wavelet transform. The period was 1.02 days (closest possible to 24 hours) because of the way the algorithm partitions the periods.

**Figure 2.**
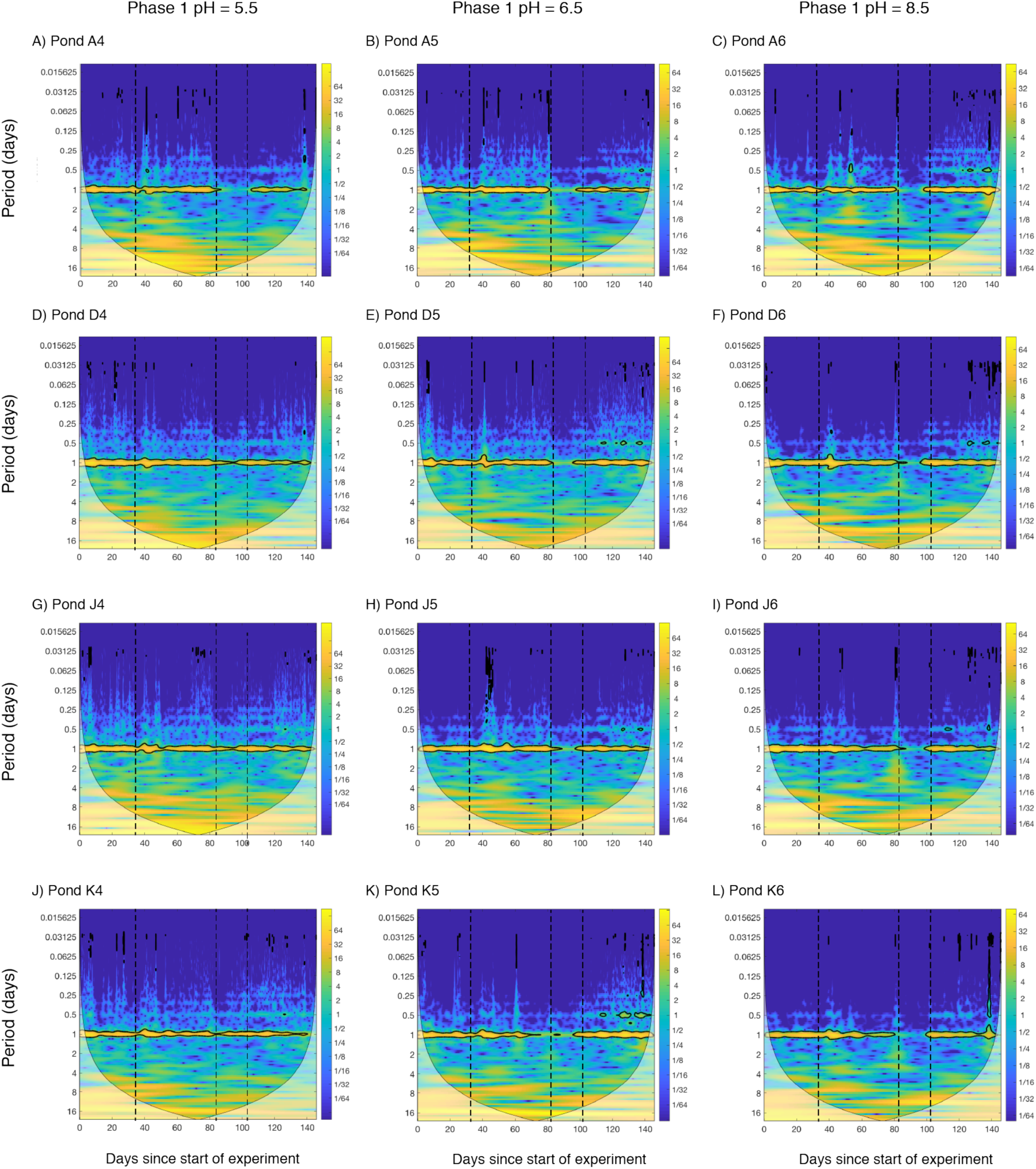
Magnitude scalograms for all ponds with Morlet wavelet. Left row, ponds at pH 5.5, middle row, ponds at pH 6.5 and right row, ponds at pH 8.5 during phase 1. Dashed lines indicate the start of phase 1 (day 34), the start of phase 2.2 (day 83) and the start of phase 2.2 (day 103). Yellow colors show high energies (“high variation”) and blue colors show low energies (“low variation”). Significant variances are circled in black. The time scale increases vertically. The cone depicts the significant area, values outside are based on too little data. See Fig. S5 for visualization of the 1-day period.

**Figure 3.**
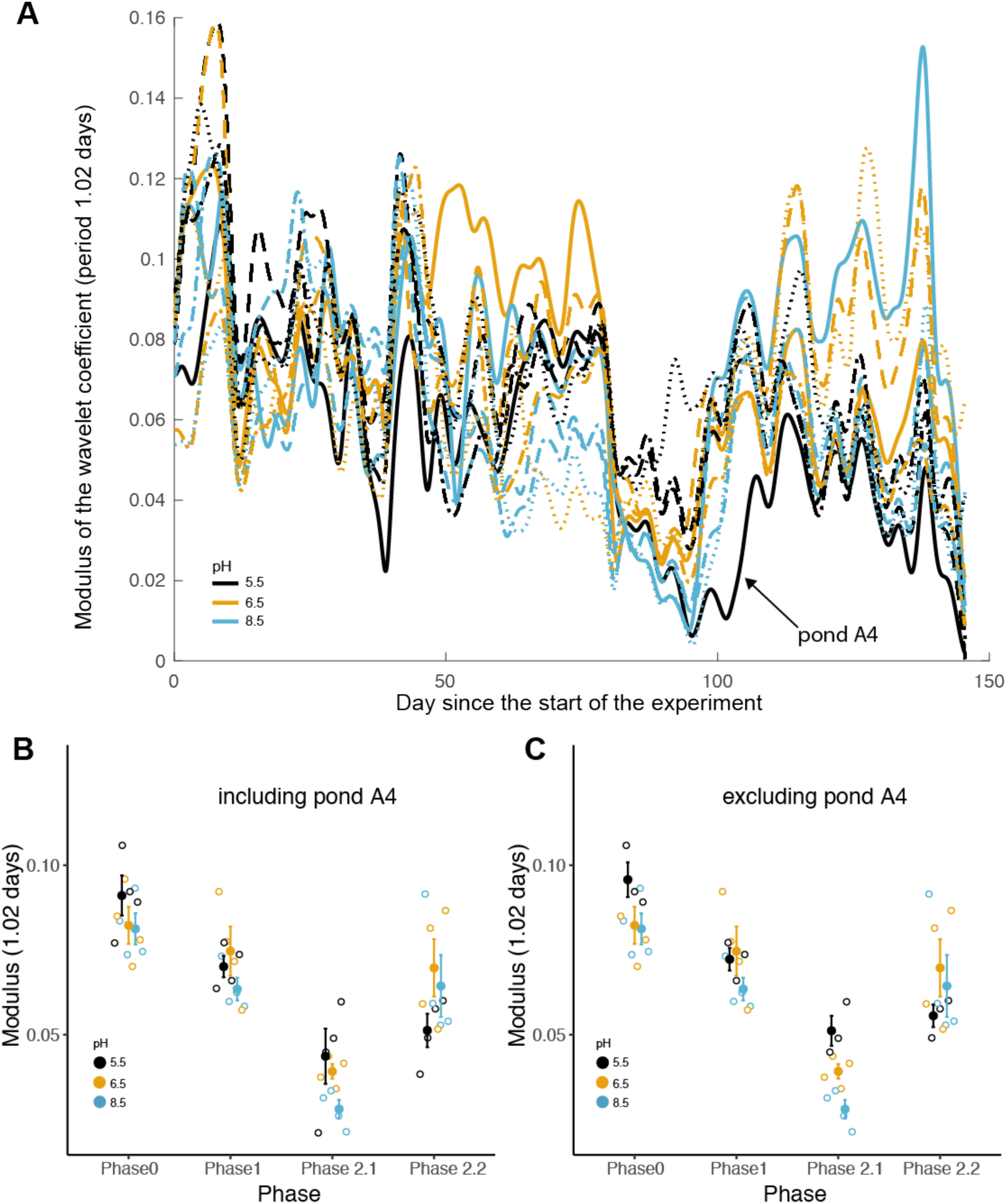
**(A)** The modulus (absolute value) of the complex coefficients for the Morse wavelet at the 1.02-period scale for each pond over time. The modulus acts like an ‘amplitude envelope’ capturing the real part of the wavelet coefficient (see Fig. S7) and describes the magnitude of the daily variation across a one-day window. **(B)** and **(C)** show means and the associated standard error for the moduli for each phase separately. In **(B)**, the outlier pond A4 is included, in **(C)** it is excluded. Black lines and points, pH 5.5; orange lines and points, pH 6.5; blue lines and point, pH 8.5.

### Statistical analysis

We tested the hypothesis that the daily DO cycle would change with pH pre-exposure treatment over the phases of the experiment. Our expectation was that the daily DO cycles would be stronger in ponds that had been pre-exposed to lower pH values 5.5>6.5>8.5 during phase 1. The time series was split into four phases and analyzed separately for each phase: phase 0 refers to the time period when no stressor was applied, phase 1 to the time period when the stressor was applied to a subset of the ponds at sub-lethal levels. The initial response of an ecosystem to a pulse perturbation can strongly differ from its long-term response (Arnoldi et al. 2018). Therefore, we separated phase 2 into two sub-phases: phase 2.1 refers to the first approximately three weeks immediately after acidification to pH 3 (August 14 to Sept 3) and phase 2.2, which refers to the rest of the field season from Sept 4 to Oct 15.

Effects of treatments on the extracted amplitudes (modulus) from the wavelet transform at a period of 1.02 days were analyzed using mixed-effects models with pH during phase 1 as a fixed-effect term (pH 5.5 *vs.* pH 6.5 *vs.* pH 8.5) and pond (n=12) as random-effect term. Mixed models using residual maximum likelihood (REML) were fitted using the package nlme for R (Pinheiro et al. 2019). All statistical analyses were conducted using the software R version 3.6.1 (R Development Core Team 2019).

## RESULTS

DO in % saturation was on average 105 % for the pH treatment 5.5 during phase 1, whereas the DO in pH treatment 6.5 was on average 117 % and thus comparable to the control treatment (pH 8.5) at 118 % saturation (Fig. 1, Table S2, Fig. S4). Upon acidification to pH 3, however, DO values were significantly higher for the two pH treatments that experienced a lower pH during phase 1 for both DO in % saturation and DO in mg/l (phase 2.1, *P* = 0.032 and *P* = 0.025, respectively, Table S2, Fig. S4).

Irrespective of the pH treatment during the pre-selective phase (phase 1), all ponds demonstrated significant daily cycles in DO (yellow band surrounded by black lines at period of 1 day, Fig. 2). Following acidification to pH 3, those ponds pre-exposed to pH 6.5 (a very mild acidic environment), or not pre-exposed to acid at all, lost their daily DO cycle for approximately two weeks (Fig. 2). In contrast, after exposure to pH 3, three out of the four ponds previously exposed to pH 5.5 during phase 1 maintained their daily DO cycle. The exception was pond “A4”, which was accidentally acidified to pH 2.5 and therefore lost its daily cycle, thus showing response more comparable with that of ponds either pre-exposed to pH 6 or not pre-exposed to an acidic environment at all. Ponds pre-exposed to pH 5.5 showed greater variation over the 24 hours after acidification to pH 3 (phase 2.1, Fig. 3B), however the effect was only significant when the outlier pond A4 was excluded (Fig. 3C, Table S1).

At the end of the experiment (phase 2.2 in Fig. 1) we observed the recovery of the daily DO cycle in all twelve ponds.

## DISCUSSION

DO is one of the most important indicators of the functioning of aquatic systems (Rajwa-Kuligiewicz et al. 2015). The amount of dissolved oxygen in the water, and in particular, the diurnal cycle of DO are measures of ecosystem metabolism and ecosystem health (Venkiteswaran et al. 2008, Demars et al. 2015, Schindler et al. 2017). We found that ponds which were pre-adapted to pH 5.5 maintained their natural DO cycle throughout the acidification, whereas ponds held at 6.5 and 8.5 did not. One might even say that the diurnal cycle of dissolved oxygen is the “heart-beat” of aquatic ecosystems. The observed crash of diurnal variability in the mesocosms held at pH 6.5 during phase 1 is thus indicative of the effect of the stress on the maintenance of ecosystem functioning. Acidification can have negative consequences for aquatic life, either via direct physiological impacts on organisms (Schindler 1985), selection driven mortality, and via changes in species abundances, and community diversity and composition (Geelen and Leuven 1986, Locke and Sprules 1994).

Despite being exposed to severe stress, we observed a recovery of the DO cycle, at the end of the experiment (phase 2.2 in Fig. 1), although the amplitudes and pattern of fluctuations differed among ponds. Because the DO cycle is mainly driven by phytoplankton (Smith and Piedrahita 1988), we also measured total algal community biomass over time, albeit at a much lower temporal resolution than DO. Acidification generally has a strong effect on algal community composition and can reduce species diversity, but at the same time it was shown that community biomass is usually little effect (Geelen and Leuven 1986). We found that the loss of the DO cycle was accompanied by a temporary and strong reduction in algal biomass (Fig. S5) and that the recovery of the DO cycle was paralleled by a recovery of algal biomass during phase 2.1. It is possible that a pure ecological process via sorting could have led to a loss of diversity, but rapid growth of the remaining tolerant algal species quickly compensated the loss of species (Klug et al. 2000). Alternatively, species may have rapidly adapted through phenotypic plasticity (Chevin et al. 2013) or evolution, either via selection on standing genetic variation (Barrett and Schluter 2008) or *de novo* mutations. Thus, preselection to stress may allow a degree of community tolerance, through a combination of ecological sorting and evolutionary selection which can also maintain ecosystem function (Bell et al. 2019, Fugère et al. 2020). Phytoplankton typically double population sizes within one to a few days, depending on the environmental conditions (Reynolds 1984). It is thus conceivable that selection pressures had an immediate evolutionary impact on the community which resulted in the DO cycle being restored.

We showed very weak effects of pre-exposure to pH 6. In part this might be because the buffering capacity of the ponds resulted in an actual pH close to 6.5. A pH of 6.5 is circumneutral and commonly experienced in Lac Hertel, the source for our ponds (Kalff 1972). Therefore, a pH of 6.5 was likely insufficient to induce a community wide tolerance to acidification. In contrast, acidification to pH 5.5 is known to reduce phytoplankton diversity and change phytoplankton community composition (Geelen and Leuven 1986) and likely constituted a stronger selective condition.

Explicit consideration of time scale is critical to modeling associations between variables measured because patterns can change both qualitatively and quantitatively with the scale of analysis (Keitt and Urban 2005, Keitt and Fischer 2006). The loss of diurnal variability represents a loss of function for these aquatic ecosystems, but over longer time scales the maintenance of cycle stability is an indication of community resistance. The wavelet analysis revealed the temporal scales at which acidification had the strongest impact. As per our hypothesis, pH strongly influenced daily fluctuations in dissolved oxygen. A coarser dataset (e.g. daily or weekly averages of DO) would have obscured our key finding. An important general conclusion is that the analysis of community and ecosystem stability requires the measurement of dynamics at multiple temporal scales, and at frequencies that can detect changes expected at the shortest scales.

### Conclusions

The rapid and severe ecological impacts associated with the human-caused contamination of aquatic ecosystems make it ever more important to study the conditions allowing communities persist and recover (Bell and Gonzalez 2011, Vander Wal et al. 2012, Geerts et al. 2015). We found that the temporal stability of the DO cycle could be maintained when exposed to extreme stress if the ponds were pre-exposed to intermediate stress (MacGillivray et al. 1995, Flöder and Hillebrand 2012, Wright et al. 2015). Prior exposure to acid stress can attenuated the impact of strong acidification on the DO cycle’s persistence and amplitude. Given the strength of the responses we hypothesize that the persistence of the DO cycle involved a joint ecological and evolutionary sorting of the phytoplankton community resulting in an acid tolerant community capable of maintaining a normal ecosystem cycle even at pH 3. More work is required to uncover the contribution of ecological and evolutionary processes to rapid adaptation and to quantify their ability to restore ecosystem stability in a stressful environment.

## Data availability

All data presented, and all code used will be archived on an online repository upon manuscript acceptance.

## Acknowledgements

The Canadian Foundation for Innovation and the Liber Ero Chair in Biodiversity Conservation provided funding to A.G. to construct the LEAP pond facility. S.J.V.M. was funded by the Swiss National Science Foundation (grant number P2ZHP3_L81462). O.L.P and S.J.V.M. would like to thank the URPP GCB of the University of Zurich for financial support. We are very grateful to Jihane Benbahtane, Rachel Rolland, Aseel Shakra, Kaushar Kagzi for data collection and David Maneli for technical support in the field.

## Author contributions

A.G., O.L.P, & S.J.V.M designed the study, S.J.V.M., C.B. & J.O.N.D. carried out the experiment, S.J.V.M, J.M. & V.F. analyzed data, S.J.V.M. wrote the paper with substantial contributions of all authors

