## Supplemental Information for "Prior selection prevents the loss of an ecosystem cycle during acidification"

***Table of Contents***


| | |
|---|---|
| Supplementary Methods | pp. 2,3 |
| Figure S1: Timeline | p. 4 |
| Figure S2: pH of the ponds | p. 5 |
| Figure S3: Pond depth | p. 6 |
| Figure S4: Mean DO in mg/l and % saturation | p. 7 |
| Figure S5: Total chlorophyll a | p. 8 |
| Figure S6: Strength of daily oscillations | p. 9 |
| Figure S7: Visualization of the modulus | p. 10 |
| Table S1: ANOVA test statistics for the (modulus) for DO | p. 11 |
| Table S2: ANOVA test statistics for mean DO | p. 12 |

***Supplementary Methods***

*Addition of* H. anomala *and* D. pulex

On June 19, we added 4 individuals of an invasive shrimp (*Hemimysis anomala*) to ½ of the ponds, i.e. in the six ponds belonging to subarrays A and K. The shrimps were collected from the St. Lawrence River at the Old Port in Montreal, QC. We did not differentiate between male/female shrimps but avoided small juveniles. Most shrimps we used were approximately 10 mm long. There were very few large individuals. To provide a refuge for the shrimps, we added to each pond a 30 cm piece of black PVC tubing (diameter of 40 mm) by weighing it down with pebbles and sinking it to the bottom of the ponds. After the addition of the shrimps, the ponds had another week to settle. In the first week of July, the south of Quebec experienced a record heat wave and outside temperatures were close to 40° C for several days in a row (Environment Canada). This resulted in elevated water temperatures (max. 38° C, see Figure 1). *H. anomala* tolerates water temperatures between 2° C and 28° C (Ricciardi et al. 2012). Therefore, the water temperatures in the pond led to 100% mortality of all shrimps previously added to the ponds. We did not find any effect of the shrimp on total chlorophyll or zooplankton counts (data not shown). Therefore, we do not further discuss this treatment in this paper.

Further, on June 5, we added 100 *Daphnia pulex* individuals to each pond as part of the full experiment, but the high water temperatures described above also resulted in very high mortality of these *D. pulex* individuals.

*Topping up of ponds*

The high air temperatures and low precipitation in early July resulted in the ponds experiencing water losses due to evaporation. To keep water levels at the desired amount of ~1000 liters, we topped up the ponds using well water. The well water originated from within the Gault Nature Reserve (about 500 m away from the field site) and was piped directly into the ponds. This well water was devoid of any algae and/or zooplankton, but still constituted a major invasion event by bacterial communities. On 4, 5 and 6 July, we added the total amount 180 liters of well water (~10% of the total volume) to the ponds. For pond depth over the field season see Figure S3.

*Wavelet analysis*

The continuous wavelet transform works by cross-correlating a time series data with a wavelet (or convoluting with the time-reversed wavelet), which is a finite, time-localized oscillation such that is non-zero only between times $t_A$ and $t_B$. The cross-correlation process works by centering the wavelet at a time t and integrating it with the data, which outputs a coefficient that describes how well the wavelet with a specific frequency (or period) matches up with the underlying data. The algorithm then expands on this by either shifting the time t or by changing the frequency up or down. Therefore, the continuous wavelet transform generates a matrix of coefficients describing the relative presence of oscillations of a specific frequency at a given time, which can be visualized in a scalogram with axes of period (or frequency) and time. For Figure 2, we utilized a Morlet mother wavelet with wavenumber 15 in order isolate frequencies at the expense of time location and to statistically test the significance of the oscillations. For Figures 3, S6 and S7 we used a Morse mother wavelet with symmetry parameter gamma equal to 3 and the time-bandwidth of 60.

*Supplementary Figures and Tables*

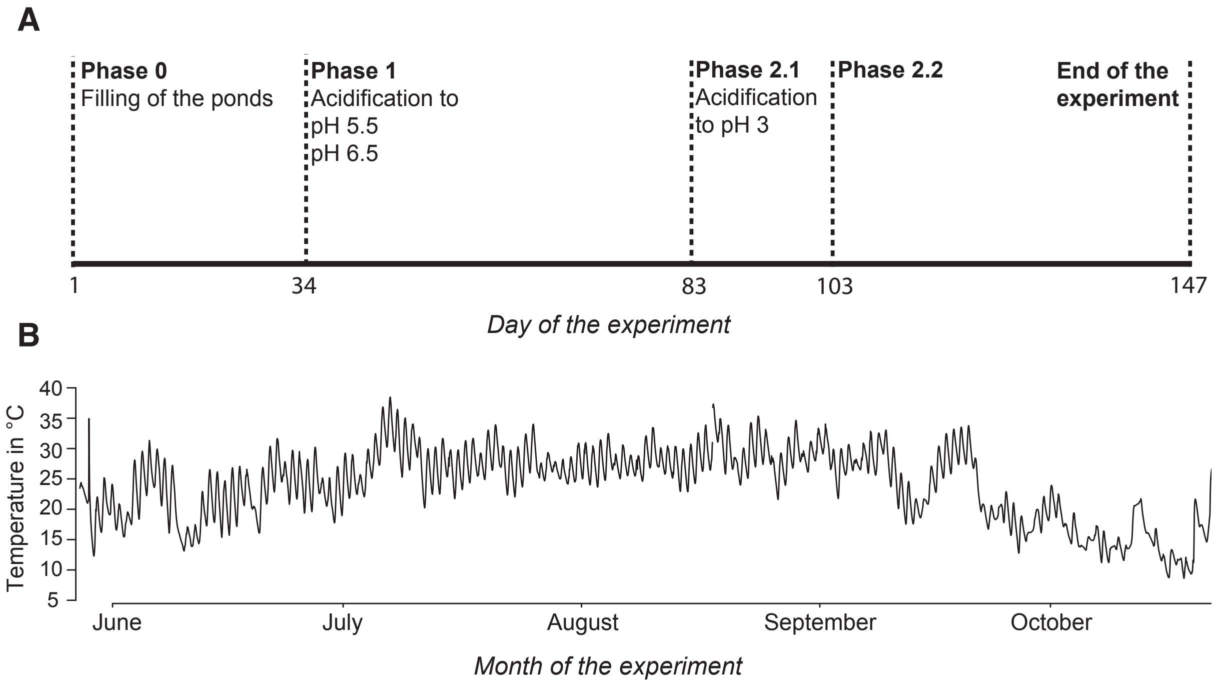


***Figure S1***. (**A**) Timeline of the experiment with the treatment manipulations indicated with a dashed line. (**B**) Water temperatures across 100 ponds over the entire field season measured by HOBO data loggers deployed in all ponds.

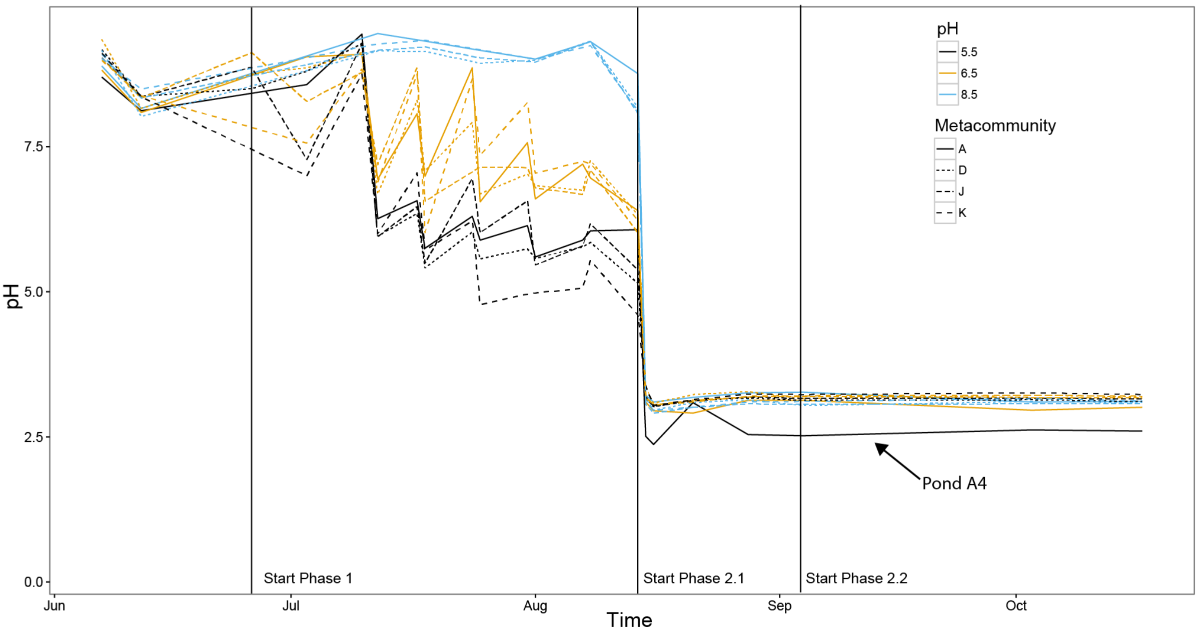


***Figure S2.*** pH in the 12 ponds with miniDOTs deployed. Metacommunity refers to the specific metacommunity (without dispersal) consisting of three ponds. Black lines, pH 5.5, yellow lines, pH 6.5, blue lines, pH 8.5. Note that pond A4 was accidentally acidified to pH 2.5, which had a strong effect on our results.

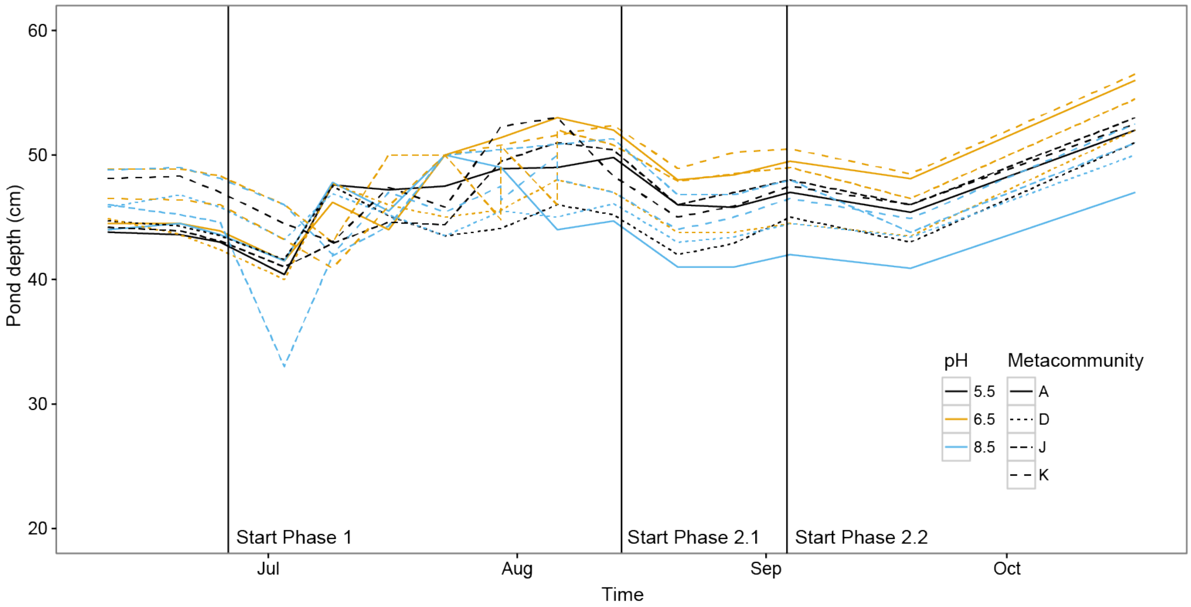


***Figure S3.*** Pond depth (in cm) for the 12 ponds with miniDOTs deployed. In July and August, ponds were topped up with well water to keep water levels around 50 cm. One particular pond (J6) had a momentarily very low water level (33cm) on July 3rd. In September and October, water levels increased due to rain fall. Black lines, pH 5.5, yellow lines, pH 6.5, blue lines, pH 8.5.

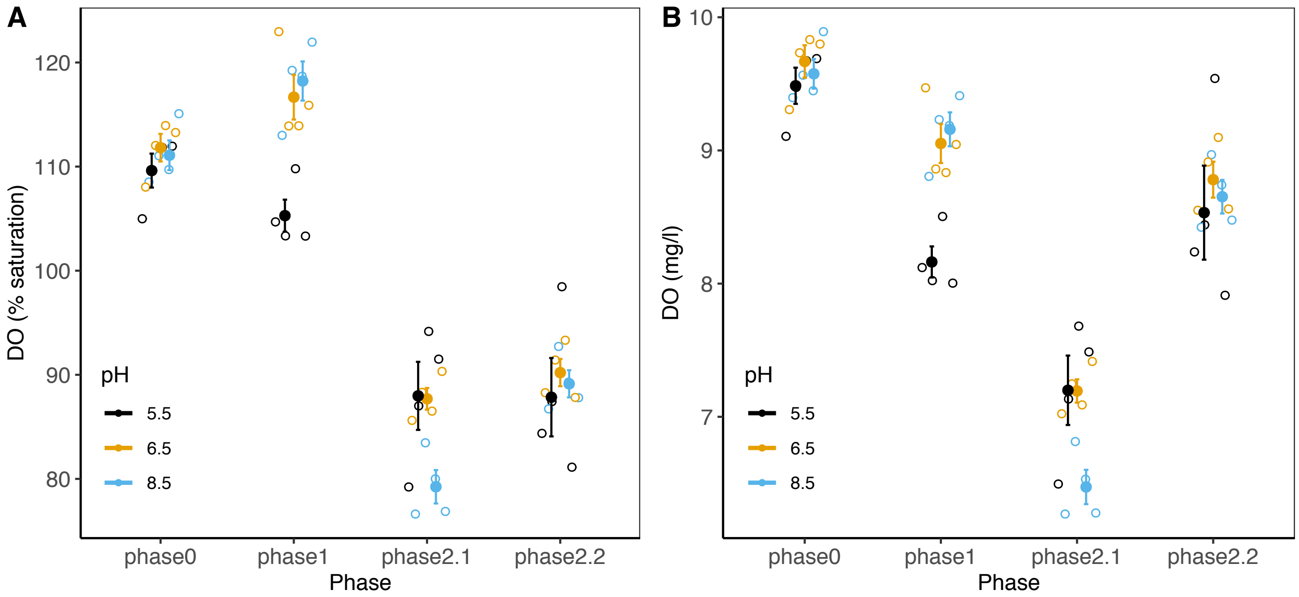


***Figure S4.*** Mean DO (**A**) in % air saturation and (**B**) mg per liter per pH treatment and phase of the experiment. During phase 1, DO for the pH 5.5 treatment was significantly lower ($P$ = 0.0066 for % saturation and $P$ = 0.021 for mg/l). Large points are means and associated standard errors (n=4 per pH treatment and phase). Note the y-axis scaling. For test-statistics see Table S1. Black points, pH 5.5, orange points, pH 6.5 and blue points, pH 8.5.

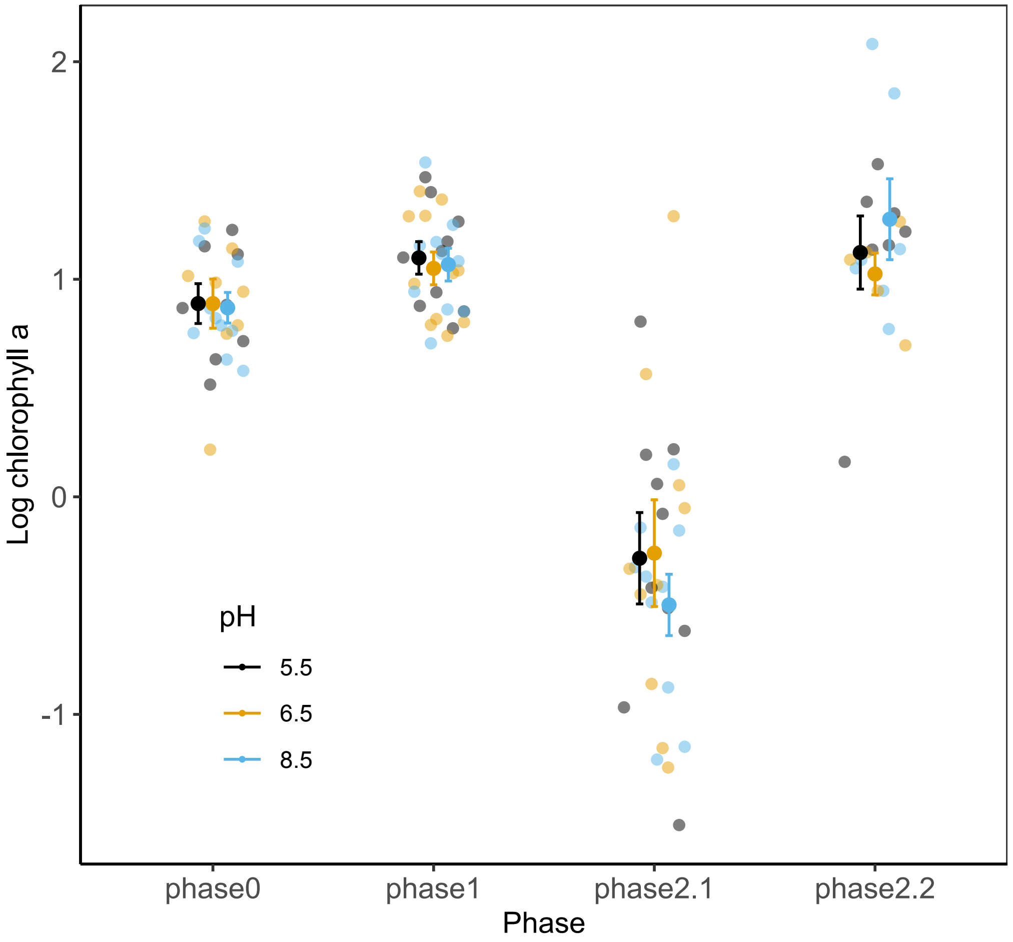


***Figure S5.*** Log-transformed total chlorophyll a (in µg/l) for the 12 ponds with miniDOTs deployed. Chlorophyll a concentration was determined fluorometrically with a FluoroProbe (bbe Moldaenke, Schwentinental, Germany). The FluoroProbe determines both total phytoplankton biomass (pigment concentration) and the biomass of four major groups: green algae (chlorophytes), golden/brown algae (diatoms, chrysophytes, and dinoflagellates), blue-green algae (cyanobacteria), and cryptophytes. Shown are means and standard errors for each phase of log-transformed total chlorophyll a concentration (darker filled circles, a proxy for total algal biomass). Each sampling point is plotted in the back (lighter circles). First sampling was on June 6, last sampling was on September 19, 2018. Data points above 60 µg/ l (n=10) were removed because they were attributed to measurement error.

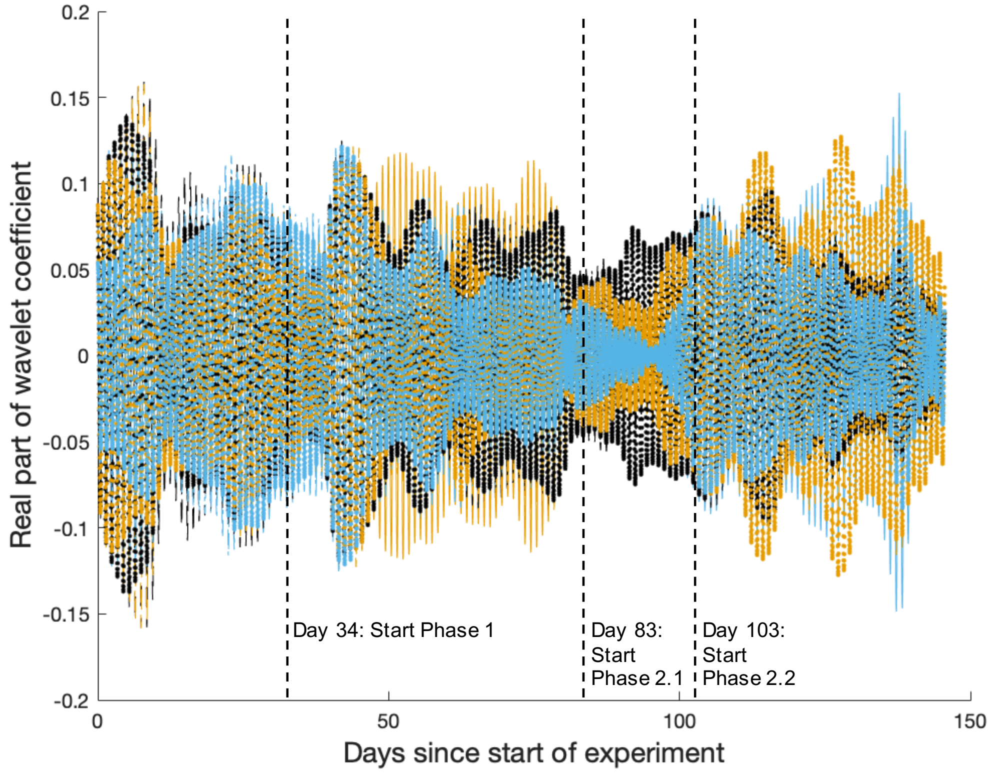


***Fig. S6***. Strength of daily oscillations in dissolved oxygen saturation for communities in ponds with pH 5.5 (black lines), pH 6.5 (yellow lines) or pH 8.5 (blue lines) before acidification to pH 3 on day 83 of the experiment. The magnitude of the real part of a Morse wavelet is used to quantify the strength of the oscillation as a zero value would indicate non-oscillatory data at the period of approximately a day (1.02 days).

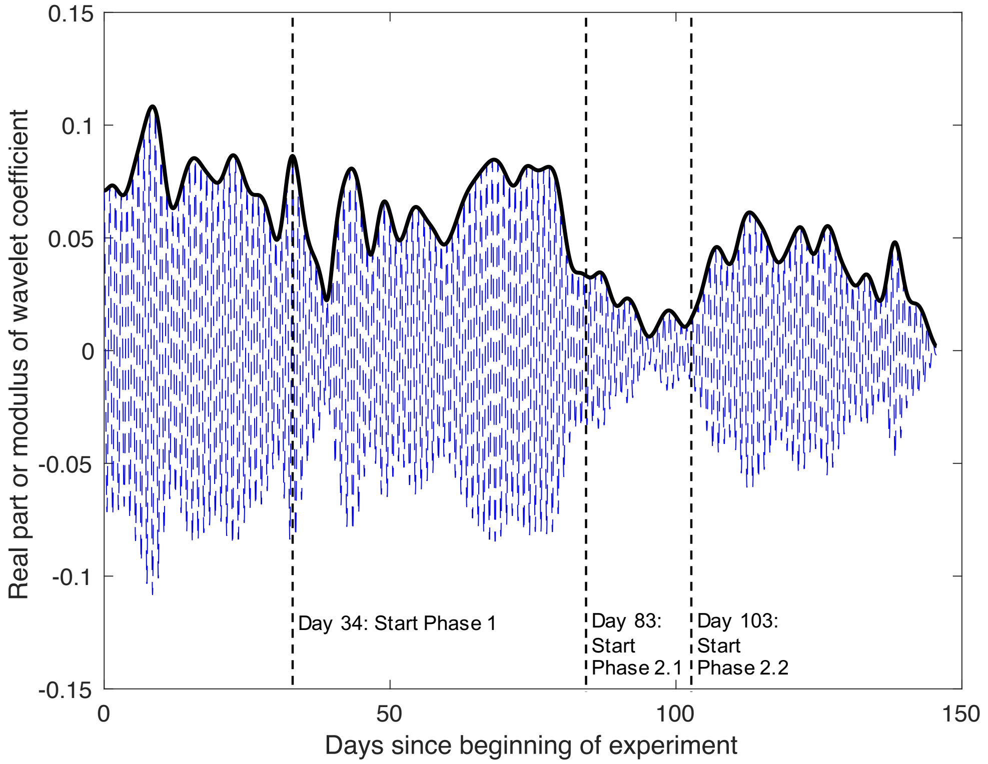


***Fig. S7.*** Visualization of the modulus for a Morse wavelet capturing the real part of the wavelet coefficient (for one example pond). Moduli for all ponds are shown in Figure 3. The modulus can be thought of as the amplitude envelope for the oscillations of the time series at the given frequency, though this only holds due to the use of the L1 norm.

**Table S1.** Results from a mixed-effects model for the variation (modulus) for DO (in % saturation) for each phase separately. The fixed-effect terms were pH and time as continuous variable, the random-effect term was pond (n=12). Significant *P*-values are shown in bold.

| *Phase* | *Including pond A4* | | | | | *Excluding pond A4* | | | |
|---|---|---|---|---|---|---|---|---|---|
| *Phase 0* | Fixed terms | DF | denDF | *F* | *P* | DF | denDF | *F* | *P* |
| | Time | 1 | 28343 | 3718.61 | **<.0001** | 1 | 25981 | 836.3 | **<.0001** |
| | pH | 2 | 9 | 1.020 | 0.3989 | 2 | 8 | 2.228 | 0.1702 |
| | Random terms | *N* | StDev | | | *N* | StDev | | |
| | Pond | 12 | 0.0073 | | | 11 | 0.01 | | |
| | Residuals | 27527 | 0.0176 | | | 25993 | 0.0192 | | |
| *Phase 1* | Fixed terms | DF | denDF | *F* | *P* | *DF* | denDF | *F* | *P* |
| | Time | 1 | 41805 | 2.32 | **<.0001** | 1 | 39060 | 4837.0 | **<.0001** |
| | pH | 2 | 9 | 1.293 | 0.3209 | 2 | 8 | 1.321 | 0.3193 |
| | Random terms | *N* | StDev | | | *N* | StDev | | |
| | Pond | 12 | 0.0074 | | | 11 | 0.0102 | | |
| | Residuals | 41818 | 0.0174 | | | 39072 | 0.0180 | | |
| *Phase 2.1* | Fixed terms | DF | denDF | *F* | *P* | DF | denDF | *F* | *P* |
| | Time | 1 | 17267 | 340.57 | **<.0001** | 1 | 15828 | 3558.5 | **<.0001** |
| | pH | 2 | 9 | 1.34 | 0.1401 | 2 | 8 | 14.078 | **0.0024** |
| | Random terms | *N* | StDev | | | *N* | StDev | | |
| | Pond | 12 | 0.0070 | | | 11 | 0.0057 | | |
| | Residuals | 17280 | 0.0133 | | | 15840 | 0.0111 | | |
| *Phase 2.2* | Fixed terms | DF | denDF | *F* | *P* | DF | denDF | *F* | *P* |
| | Time | 1 | 32140 | 367.93 | **<.0001** | 1 | 34374 | 10341.7 | **<.0001** |
| | pH | 2 | 9 | 1.00 | 0.2738 | 2 | 8 | 0.709 | 0.5205 |
| | Random terms | *N* | StDev | | | *N* | StDev | | |
| | Pond | 12 | 0.0158 | | | 11 | 0.0156 | | |
| | Residuals | 32153 | 0.0198 | | | 34386 | 0.0147 | | |

**Table S2.** Results from a mixed-effects model for pH treatment on DO (in % saturation) for each phase separately. The fixed-effect terms were pH and time as continuous variable, the random-effect term was logger (n=12). Significant P-values are shown in bold.

| *Phase* | | *in % saturation* | | | |
|---|---|---|---|---|---|
| *Phase 0* | Fixed terms | DF | denDF | *F* | *P* |
| | Time | 1 | 27514 | 3153.13 | **<.0001** |
| | pH | 2 | 9 | 0.58 | 0.579 |
| | Random terms | *N* | StDev | | |
| | logger Id | 12 | 2.93 | | |
| | Residuals | 27527 | 11.626 | | |
| *Phase 1* | Fixed terms | DF | denDF | *F* | *P* |
| | Time | 1 | 41805 | 16923.69 | **<.0001** |
| | pH | 2 | 9 | 14.20 | **0.002** |
| | Random terms | *N* | StDev | | |
| | logger Id | 12 | 3.74 | | |
| | Residuals | 41818 | 9.247 | | |
| *Phase 2.1* | Fixed terms | DF | denDF | *F* | *P* |
| | Time | 1 | 17267 | 20.69 | **<.0001** |
| | pH | 2 | 9 | 5.15 | **0.032** |
| | Random terms | *N* | StDev | | |
| | logger Id | 12 | 4.37 | | |
| | Residuals | 17280 | 8.699 | | |
| *Phase 2.2* | Fixed terms | DF | denDF | *F* | *P* |
| | Time | 1 | 32140 | 2954.64 | **<.0001** |
| | pH | 2 | 9 | 0.22 | 0.808 |
| | Random terms | *N* | StDev | | |
| | logger Id | 12 | 4.84 | | |
| | Residuals | 32153 | 7.482 | | |

**Note:** *DF, degrees of freedom, denDF, denominator degrees of freedom*
